# Context-Aware Seeds for Read Mapping

**DOI:** 10.1101/643072

**Authors:** Hongyi Xin, Mingfu Shao, Carl Kingsford

**Affiliations:** Computer Science Department, School of Computer Science, Carnegie Mellon University, Pittsburgh, PA; Department of Computer Science and Engineering, The Pennsylvania State University, University Park, PA; Computational Biology Department, School of Computer Science, Carnegie Mellon University, Pittsburgh, PA

## Abstract

**Motivation:** Most modern seed-and-extend NGS read mappers employ a seeding scheme that requires extracting *t* non-overlapping seeds in each read in order to find all valid mappings under an edit distance threshold of *t*. As *t* grows (such as in long reads with high error rate), this seeding scheme forces mappers to use more and shorter seeds, which increases the seed hits (seed frequencies) and therefore reduces the efficiency of mappers.

**Results:** We propose a novel seeding framework, context-aware seeds (CAS). CAS guarantees finding all valid mapping but uses fewer (and longer) seeds, which reduces seed frequencies and increases efficiency of mappers. CAS achieves this improvement by attaching a confidence radius to each seed. We prove that all valid mappings can be found if the sum of confidence radii of seeds are greater than *t*. CAS generalizes the existing pigeonhole-principle-based seeding scheme in which this confidence radius is implicitly always 1. Moreover, we design an efficient algorithm that constructs the confidence radius database in linear time. We experiment CAS with *E. coli* genome and show that CAS reduces seed frequencies by up to 25.4% when compared with the state-of-the-art pigeonhole-principle-based seeding algorithm, the Optimal Seed Solver.

**Availability:** https://github.com/Kingsford-Group/CAS_code

## 1 Introduction

Read mapping is used ubiquitously in bioinformatics. Commonly, it is defined as follows:

### Problem 1

(Read Mapping). *Given read R and reference T (usually with |T | ≫ |R|), an edit distance measurement D*(*·, ·*), *and an error tolerance threshold t, we say a substring of T at location* [*l*_1_, *l*_2_], *i.e., T* [*l*_1_, *l*_2_], *is a valid mapping of R if we have D*(*R, T* [*l*_1_, *l*_2_]) *< t.*

To efficiently map reads, modern mappers usually employ the *seed-and-extend* mapping strategy [1, 8, 9, 14]: a mapper extracts a substring of *R* as a *seed, s*; iterates through all seed locations of *s* in *T*; at each seed location, performs sequence alignment of *R* against the surrounding text in *T*; reports alignments that have edit distances below *t* as valid mappings.

For mappers that use non-overlapping seeds, the number of seeds to extract from a read *R* is governed by the pigeonhole principle: to find all valid mappings of *R*, the mapper must divide *R* into at least *t* non-overlapping seeds. Otherwise, the mapper will not be able to consistently find all valid mappings of *R* in *T*. As *t* grows, the length of seeds is reduced. Using short seeds significantly increases the workload of a mapper [6, 11]. Shorter seeds appear more frequently in *T,* hence increaseing the number of alignments while mapping a read. To improve the performance of mappers, it is desirable to use fewer non-overlapping seeds under a fixed *t*, which lets a mapper not only use fewer seeds, but also use longer seeds.

In this paper, we focus on improving seed-and-extend mappers that use non-overlapping seeds. We propose a novel seeding scheme, called *context-aware* seeds (CAS). CAS enables a mapper to use fewer than *t* seeds without missing any valid mappings. CAS attaches each seed *s* with a *confidence radius* score, *c*_*s*_, with *c*_*s*_ *≥* 1. Let *S* be a set of non-overlapping seeds from *R*. CAS ensures that as long as Σ_*s∈S*_ *c*_*s*_ *≥ t*, then *S* is sufficient to find all valid mappings of *R* under an error tolerance threshold of *t*. When *S* includes any seed *s* with *c*_*s*_ *>* 1, then *|S| < t* and all valid mappings are secured with fewer-than-*t* seeds (*|S|* denotes the number of seeds in *S*). In the worst case where *c*_*s*_ = 1 for all *s ∈ S*, CAS degenerates into the case governed by pigeonhole principle with *|S|* = *t*.

Figure 1 compares CAS and the pigeonhole-principle-based seeds. Assume that we have verified that the two CAS seeds `AACC` and `TTGG` have confidence radii of *c*_*s*_ = 2. Therefore CAS can be guaranteed to find all valid mappings with just these two seeds, as Σ_*s*_ *c*_*s*_ = 4 ≥ *t*. Using the pigeonhole principle, however, a mapper needs to select *t* = 4 non-overlapping seeds. It forces the mapper to pick short and repetitive seeds, making the mapper perform more local alignments.

**Figure 1:**
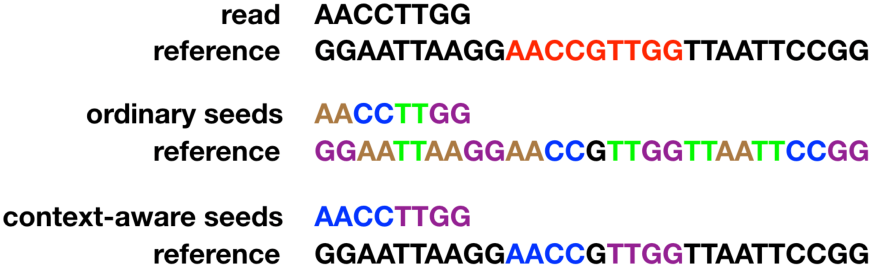
Illustration of CAS. The upper part shows a read and a reference. Suppose that *t* = 4, i.e., we want to find all alignments of the read in the reference with fewer than 4 edits. There is only one such locally optimal alignment (marked as red). The middle part shows the seed extraction result with the pigeonhole principle, which splits the read into *t* = 4 seeds. This gives many seed locations and thus many alignments. With CAS (in the lower part), we can split the read into 2 long seeds while still guarantee to find all valid mappings. The two long seeds together have a total seed frequency of 2, drastically reducing the number of alignments.

We establish the theoretical foundation of CAS and demonstrate that with CAS future mappers can map reads more efficiently using fewer, longer and less frequent seeds without losing valid mappings. We also propose a suffix-trie-based CAS database construction algorithm that builds a CAS database from *T* in linear time, based on which we design a greedy CAS seeding algorithm that extracts CAS from reads. We test the greedy CAS seeding algorithm against a state-of-the-art pigeonhole-principle-based seeding algorithm, Optimal Seed Solver (OSS), on an *E. coli* dataset.

## 2 Context-Aware Seeds

Let *s* be a string and [*l*_1_, *l*_2_] be a pair of locations. We say string *T* [*l*_1_, *l*_2_] is in the *vicinity* of *s* under *t*, if *∃*[*l*_*s*1_, *l*_*s*2_], where *l*_1_ *- t < l*_*s*1_ *< l*_*s*2_ *< l*_2_ + *t* and *T* [*l*_*s*1_, *l*_*s*2_] = *s*. Furthermore, let seed *s* be a substring of *R* at [*l*_*r*1_, *l*_*r*2_] (*s* = *R*[*l*_*r*1_, *l*_*r*2_]) and let *T* [*l*_1_, *l*_2_] be a valid mapping of *R*. We say *T* [*l*_1_, *l*_2_] is in the *vicinity of s with regard to R* under *t*, if string 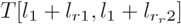 is in the vicinity of *s* under *t*. If a valid mapping *T* [*l*_1_, *l*_2_] is in the vicinity of *s* with respect to *R*, then *T* [*l*_1_, *l*_2_] can be discovered by locally aligning *R* against the surrounding text in *T* at each seed location of *s*.

The pigeonhole principle states that by dividing *R* into a set of *t* non-overlapping seeds, denoted by *S*, then *∀* [*l*_1_, *l*_2_] where *T* [*l*_1_, *l*_2_] is a valid mapping of *R*, there must be *s ∈ S* where *T* [*l*_1_, *l*_2_] is in the vicinity of *s* with regard to *R*.

CAS seeks to retain the seed vicinity guarantee of the pigeonhole principle, where all valid mappings of a read *R* are in the vicinity of its seeds with regard to *R* under *t*, with **fewer than** *t* **seeds**. Given two substrings *s* and *s′* of *T* and a edit-distance threshold *t*, we say *s′* is a neighbor of *s* if *D*(*s, s′*) *< t*. Assume that *s′* is a neighbor of *s* under *t*, CAS defines *s′* as a *trivial neighbor* of *s*, if and only if *∀* [*l*_1_, *l*_2_] where *T* [*l*_1_, *l*_2_] = *s′, T* [*l*_1_, *l*_2_] is in the vicinity of *s* under *t*. Otherwise CAS defines *s′* as a *nontrivial neighbor* of *s*. Finally, CAS defines *the confidence radius c*_*s*_ of *s* as the minimum of 1) *t* and 2) the minimum edit-distance between *s* and all nontrivial neighbors of *s*. Since a seed is trivial to itself and is at least 1-edit-distance away from any other string, we have *t ≥ c*_*s*_ *≥* 1 for any seed *s*.

We now give the central theorem of CAS, the theoretical foundation that enables seed-and-extend mappers to find all valid mappings using fewer than *t* seeds.

### Theorem 1.

*Let S is a set of non-overlapping seeds of a read R, if as* Σ_*s∈S*_ *c*_*s*_ *≥ t, then ∀*[*l*_1_, *l*_2_] *where D*(*R, T* [*l*_1_, *l*_2_]) *< t, ∃s ∈ S where T* [*l*_1_, *l*_2_] *is in the vicinity of s.*

*Proof.* Assume that *T* [*l*_1_*′, l*_2_*′*] is a valid mapping of *R*, where *D*(*R, T* [*l*_1_*′, l*_2_*′*]) *< t*. Further assume that *T* [*l*_1_*′, l*_2_*′*] is not in the vicinity, with regard to *R* under *t*, of any *s ∈ S*. In the minimum-edit-distance alignment between *R* and *T* [*l*_1_*′, l*_2_*′*], assume that the non-overlapping seeds *s*_1_, *s*_2_,…, *s*_*n*_ of *R* are aligned to the non-overlapping segments *s*_*T* 1_, *s*_*T* 2_,…, *s*_*T n*_ of *T* [*l*_1_*′, l*_2_*′*], with *n* = *|S|*. Since *T* [*l*_1_*′, l*_2_*′*] is not in the vicinity, with regard to *R* under *t*, of any *s ∈ S*; and also because *c*_*si*_ *≤ t* for all *i*; there does not exist *i* where *s*_*T i*_ is in the vicinity of *s*_*i*_, under *c*_*si*_. Therefore, *s*_*T i*_ is a nontrivial neighbor of *s*_*i*_ for all *i ∈* [1, *n*]. Because *c*_*si*_ is the minimum edit-distance between *s*_*i*_ and any of its nontrivial neighbors, we have *D*(*R, T* [*l*_1_*′, l*_2_*′*]) *≥* Σ_*i*_ *D*(*s*_*i*_, *s*_*T i*_) *≥* Σ_*s*_ *c*_*s*_ *≥ t. D*(*R, T* [*l*_1_*′, l*_2_*′*]) *≥ t* contradicts the assumption that *T* [*l*_1_*′, l*_2_*′*] is a valid mapping of *R*. Therefore such *T* [*l*_1_*′, l*_2_*′*] does not exist. □

## 3 Construction of Confidence Radius Database

The confidence radius *c*_*s*_ of each seed *s* is stored in a table, called *the confidence radius database*. The confidence radius database only needs to be constructed once offline for a reference *T*.

Computing *c*_*s*_ of seed *s* involves finding the minimum edit distance to its nontrivial neighbors. Below we propose an algorithm that constructs the confidence radius database in *O*(*|*Σ*|*^2^ · *M)* time, where Σ is the alphabet set of *T* and *M* is the total number of neighbors of all strings in *T* (up to length *P* and under the edit distance threshold *t*).

The confidence radius database is constructed in two steps: first, we construct a neighbor database, which stores all neighbors of all seeds (up to length *P)* under the edit distance threshold *t*; then, we find the confidence radius of each from its neighbors. We prove that both steps can be done in *O*(*|*Σ*|*^2^ · *M)* time.

### 3.1 Construction of the Neighbor Database

To find all neighbors of all substrings in *T* (up to a maximum length *P)*, we first build a *P*-level suffix trie of *T,* then find all neighbors of each seed in *Trie* by systematically traversing the suffix trie in a top-down manner. Formally, let *Trie* = (*V, E*) be a suffix trie of *T* of a maximum depth of *P* + *t*. Let *r ∈ V* be the root of *Trie*. Each node represents a substring in *T,* i.e., the string obtained by concatenating the letters on edges along the path from *r* to *v*. We denote the edit distance between these two substrings corresponding to *u* and *v* as *D*(*u, v*). We aim to solve the following problem:

#### Problem 2.

*Given a suffix trie Trie* = (*V, E*) *and an integer t, to compute all pairs of nodes u, v ∈ V such that D*(*u, v*) *≤ t.*

For any *v ∈ V, p*(*u*) denotes the parent node of *v* in *Trie. Σ*(*p*(*v*), *v*) denotes the letter on the edge between *v* and *p*(*v*), i.e., (*p*(*v*), *v*) *∈ E*. We have the following lemmas.

#### Lemma 1.

*Let u, v ∈ V. Then D*(*u, v*) *≤ t only if D*(*p*(*u*), *p*(*v*)) *≤ t.*

*Proof.* Proved in Landau and Vishkin [7] by enumerating and validating all possible scenarios. □

#### Lemma 2.

*Let u, v ∈ V. We have*

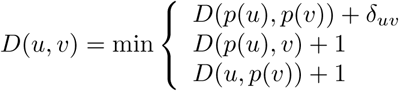

*where δ*_*uv*_ = 1 *if σ*(*p*(*u*), *u*) *≠ σ*(*p*(*v*), *v*) *and δ*_*uv*_ = 0 *if σ*(*p*(*u*), *u*) = *σ*(*p*(*v*), *v*).

*Proof.* This follows the dynamic programming algorithm for the edit distance problem. □

Lemma 1 shows that nodes are neighbors only if their parents are neighbors. Hence the neighbors of a child node must be the children of the neighbors of its parent node. Lemma 2 further shows that the edit distance between two children nodes can be computed in constant time, given the edit distances between one child and the parent node of the other child, as well as the edit distance between the two parent nodes.

We construct the neighbor database by traversing *Trie* as follows: First, assign each node in *V* an integral *rank* from {1, 2, *…,* |*V*|} following a top-down, left-to-right order. The root *r* of *Trie* has rank of 1, and then the children of children of *r* have ranks of 2, 3, from the leftmost child to the rightmost child, and so on. Nodes that are deeper in *Trie* rank higher. Among nodes of the same depth, children of a higher ranking parent node rank higher. A breadth-first-search traversal of *Trie* ranks all nodes.

For any *v ∈ V,* we define *X*_*v*_:= *{u ∈ V | D*(*u, v*) *≤ t}* as the set of neighbors of *v*, including *v* itself, and define *Y*_*v*_:= *{D*(*u, v*) *| u ∈ X*_*v*_*}* as the accompanying edit-distance set of *X*_*v*_. For every neighbor node *u* in *X*_*v*_, *Y*_*v*_ provides the edit distance between *u* and *v*. We compute *X*_*v*_ and *Y*_*v*_ for each node *v ∈ V* from low ranking nodes to high ranking nodes. Both *X*_*v*_ and *Y*_*v*_ are implemented as arrays.

The algorithm for constructing the neighbor database is summarized in Algorithm 1. We iterate through all nodes by rank from low to high. For each node *v ∈ V,* we iterate through all children of *v*. For each children node *v′* of *v*, we compute *X*_*v′*_ and *Y*_*v′*_ of *v′* based on the previously computed *X*_*v*_ and *Y*_*v*_ of *v*. Figure 2 illustrates the process of validating a candidate neighbor *u′* of another node *v′*, based on the information of its parent node *v* and the neighbor *u* of *v*, where *u* is also the parent of *u′* (line 4-17 in Algorithm 1). We prove that this algorithm maintains the following three invariants:

**Figure 2:**
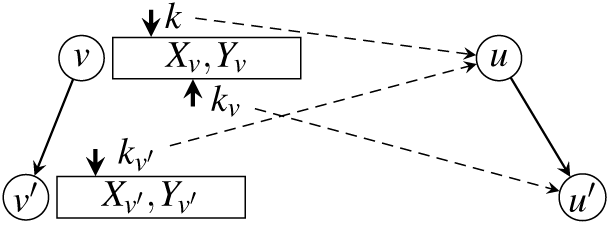
Illustration of processing a single node *v* (i.e., line 4 to line 17 of Algorithm 1).

#### Algorithm 1: Linear Time Algorithm for Problem 2

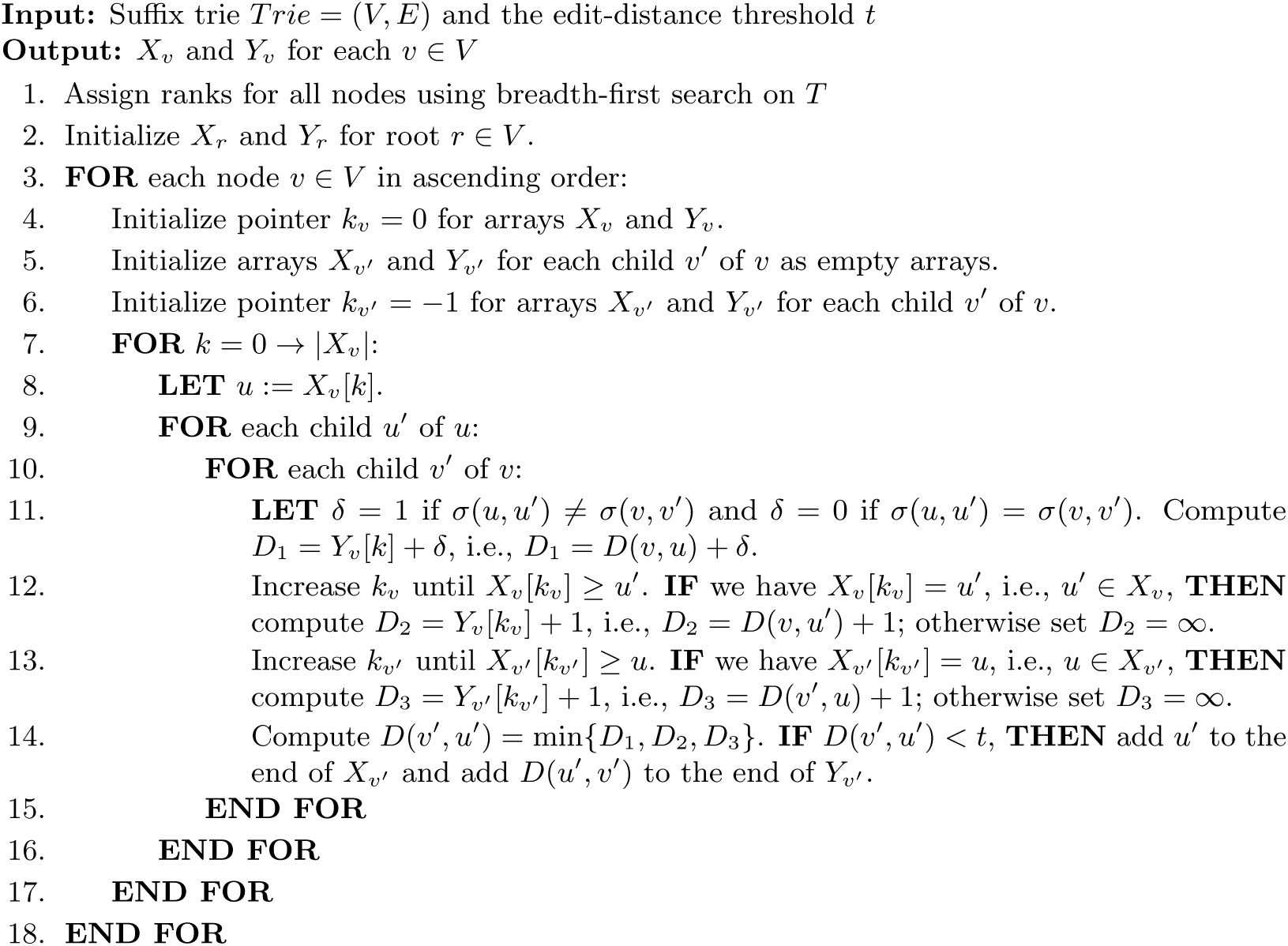

1. For any node *v ∈ V,* array *X*_*v*_ is always sorted according to their ranks, i.e., nodes that are added to *X*_*v*_ are always in ascending order *w.r.t.* their ranks.
2. Right before processing node *v* (i.e., before line 4 of Algorithm 1), *X*_*v*_ and *Y*_*v*_ are already computed and sorted *w.r.t.* their ranks.
3. Right after processing node *v* (i.e., after line 17 of Algorithm 1), *X*_*v′*_ and *Y*_*v′*_ are computed and sorted *w.r.t.* their ranks for each child *v′* of *v*.

The initialization step Algorithm 1 (line 2) computes *X*_*r*_ and *Y*_*r*_ for root node *r*. Its neighbors include all nodes whose depth in *Trie* is no greater than *t*. The edit distance between *r* to a neighbor node *u* is simply the depth of *u* minus 1 (we assume that root *r* is at depth 1). Root *r* is also in *X*_*r*_ with *D*(*r, r*) = 0. Clearly, the first and the second invariant hold for root *r*.

In the main loop (line 3-18), for a node *v ∈ V,* Algorithm 1 iterates through all of its children. For a child *v′* of *v*, line 4–17 compute *X*_*v′*_ and *Y*_*v′*_ of *v′*. Line 4-6 initialize the pointers that will be used to fetch the edit distances *D*(*v, u′*) and *D*(*v′, u*), which are stored in *Y*_*v*_ and the partially computed *Y*_*v′*_, respectively. Because *u* rank higher than *u′*, by the time of computing *D*(*v′, u′*), *D*(*v′, u*) is already computed and stored in *X*_*v′*_. *D*(*v′, u′*) is then computed according to Lemma 2. Specifically, pointer *k* tracks the position of *u* in array *X*_*v*_ (i.e., the index of *u* in array *X*_*v*_); pointer *k*_*v*_ tracks the position of *u′* in array *X*_*v*_; and pointer *k*_*v′*_ tracks the position of *u* in array *X*_*v′*_. Line 11 computes *D*_1_:= *D*(*v, u*) + *δ*, in which *D*(*v, u*) is fetched from *Y*_*v*_ indexed by *k*. Line 12 computes *D*_2_:= *D*(*v, u′*) + 1, in which *D*(*v, u′*) is fetched from *Y*_*v*_ indexed by *k*_*v*_. Line 13 computes *D*_3_:= *D*(*v′, u*) + 1, in which *D*(*v′, u*) is fetched from *Y*_*v′*_ indexed by *k*_*v′*_. Line 14 computes *D*(*v′, u′*):= min *D*_1_, *D*_2_, *D*_3_; adds *u′* to *X*_*v′*_ and adds *D*(*v′, u′*) to *Y*_*v′*_ if *D*(*v′, u′*) *< t*.

Algorithm 1 maintains the first invariant. For each child *v′* of *v*, assuming *X*_*v*_ is sorted, then neighbors are also added to *X*_*v′*_ in a sorted manner, as Algorithm 1 iterates through neighbors ordered by *X*_*v*_. Since *X*_*r*_ is sorted for root *r*, given the inductive nature of Algorithm 1, we conclude that *X*_*v*_ must be sorted for any *v ∈ Trie*.

Algorithm 1 maintains the third invariant. According to Lemma 1, a node *u′ ∈ X*_*v′*_ requires *u ∈ X*_*v*_ for their parents *u* and *v*. Any node *ū′* whose parent *ū* ∉ *X*_*v*_ results in *ū′ ∉ X*_*v′*_. Algorithm 1 iterates through all *u* in *X*_*v*_. Therefore, after line 17, all neighbors of child *v′* must have been found, assuming the second invariant holds. The second invariant holds because all neighbors of *r* are correctly defined during initialization. As the algorithm propagates, because of the inductive nature of Algorithm 1, the second invariant holds.

Let *M* denote the member size of set *{*(*u, v*) *| D*(*u, v*) *≤ t}*. The complexity of Algorithm 1 is *O*(*|*Σ*|*^2 ·^ *M)*.

#### Theorem 2.

*Algorithm 1 computes X*_*v*_ *and Y*_*v*_ *for each v ∈ V in O*(*|*Σ*|*^2^ *· M) time.*

*Proof.* For each *v* ∈ *V,* lines 4-17 compute *X*_*v′*_ and *Y*_*v′*_ for each child *v′* of *v* in *O*(|*X* Σ _*v*_ | ·^2^ + Σ _*v′v′v*_ |*X*_*v′*_ |) time. Since pointers of *k*_*v*_ and *k*_*v′*_ can only move forward, lines 12-13 cost *|X*_*v*_*|* Σ _*v′v′v*_ |*X*_*v′*_ | operations. Operations in lines 11–14 cost constant time. Hence, lines 7–17 cost *O*(*|X*_*v*_*|* · *|*Σ*|*^2^) operations, as the number of children of each node is bounded by *|*Σ*|*. The overall run time of Algorithm 1 is thus bounded by Σ_*v∈V*_ *O*(*|X*_*v*_*|*·*· |*Σ*|*^2^ + Σ _*v′:p(v′)=v*_ |*X*_*v′*_ | = *O*(*|*Σ*|*^2^ · *M)*.

With *|*Σ*|* being a small constant (for example Σ = {*A, C, G, T}* for DNA analysis), Algorithm 1 finds all *M* neighbor pairs in *Trie* in *O*(*M)* time.

### 3.2 Computing the Confidence Radius Among Nontrivial Neighbors

The neighbor database stores both the trivial and nontrivial neighbors of each seed. However, CAS requires the minimum edit distance to just the nontrivial neighbors of each seed. In order to derive the confidence radius of each seed, we propose an augmentation to Algorithm 1, such that it computes the minimum edit distance to nontrivial neighbors while constructing the neighbor database. We prove that the augmentation does not increase the time complexity of Algorithm 1.

Within the neighbor array *X*_*v*_ of a node *v*, let the sub-array 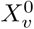 store all trivial neighbors and 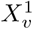 store all nontrivial neighbors, where 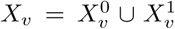. By definition, the confidence radius of *v* is computed as 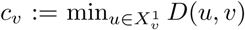. To compute *c*_*v*_, instead of finding all nontrivial neighbors, 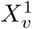, we compute a subset 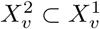, where 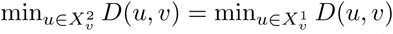.

Let *u* be a neighbor of *v*; we say *u* is an *immediate* neighbor of *v* if *u* is a substring, or a superstring, or an overlapping string of *v*; otherwise we say *u* is a *non-immediate* neighbor of *v* (see Figure 3 for examples). Immediate neighbors are not necessarily trivial neighbors. If *u* is a trivial neighbor of *v*, by definition, then *u* must be an immediate neighbor of *v*. However, the opposite is not necessarily true, i.e., *u* could be a substring of *v* (an immediate neighbor) yet *u* is nontrivial to *v*. Substring *u* may appear at more locations in *T* than *v* does. It is easier to determine whether *u* is an immediate neighbor to *v* than whether *u* is a trivial neighbor to *v*.

**Figure 3:**
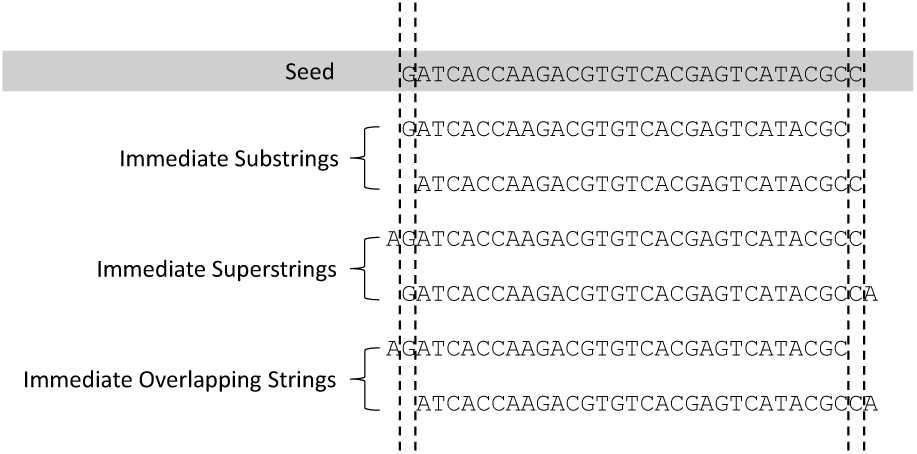
Examples of trivial neighbors of a seed, including substrings, superstrings, and overlapping strings of this seed.

Let 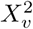 be the set of non-immediate neighbors of a node *v*. The minimum edit distance between *v* to nontrivial neighbors of *v* equals to the minimum edit distance between *v* to neighbors in 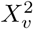. We prove this in Theorem 3. To prove Theorem 3, we first prepare the following two lemmas.

#### Lemma 3.

*If u is a superstring of v, then u is a trivial neighbor of v.*

*Proof.* Since *u* is a superstring of *v*, for any location [*l*_1_, *l*_2_] of *u, ∃*[*l*_1_, *l*_2_] where *T* [*l*_1_, *l*_2_] = *v* and *l*_1_ *- D*(*u, v*) *≤ l*_1_ *< l*_2_ *≤ l*_2_ + *D*(*u, v*). By definition, *u* is a trivial neighbor of *v*. □

#### Lemma 4.

*If u is a substring or an overlapping string of v and u is a nontrivial neighbor of v, then ∃w ∈ Trie, where w is neither an immediate neighbor nor a trivial neighbor of v, with |w|* = *|v| and D*(*v, w*) *≤ D*(*v, u*).

*Proof.* Since *u* is a nontrivial neighbor of *v, ∃*[*l*_1_, *l*_2_], where *T* [*l*_1_, *l*_2_] = *u* but *T* [*l*_1_, *l*_2_] is not in the *D*(*v, u*)-edit vicinity of *v*. We extract a substring *w* within *T* [*l*_1_ *- D*(*u, v*), *l*_2_ + *D*(*u, v*)], where *w* locally and optimally aligns to *v* in *T* [*l*_1_ *- D*(*u, v*), *l*_2_ + *D*(*u, v*)], with *|w|* = *|v|*, as shown in Figure 4. Then *w* must be a nontrivial neighbor of *v* since *T* [*l*_1_, *l*_2_] is not in the *D*(*u, v*)-edit vicinity of *v*. Because *w* is optimally aligned to *v* within [*l*_1_ *- D*(*u, v*), *l*_2_ + *D*(*u, v*)], we have *D*(*w, v*) *≤ D*(*u, v*). □

By combining Lemmas 3 and 4 we prove the following theorem.

#### Theorem 3.

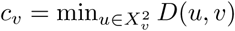, where 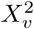 *is the set of non-immediate neighbors of v.*

*Proof.* Lemmas 3 and 4 state that for any nontrivial immediate neighbor *u* of seed *v*, there must exist a nontrivial and non-immediate neighbor *w* of *v* where *D*(*w, v*) *≤ D*(*u, v*). Therefore, by definition, we have 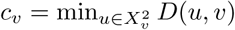. □

We find the immediate neighbors, 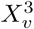, of each node *v* ∈ *Trie*, by checking if a neighbor *u* ∈ *X*_*v*_ is a immediate substring, superstring or overlapping string of *v*. We associate with each node *v* a new vector *Z*_*v*_:= *{F* (*v, u*) *| u ∈ X*_*v*_*}*, where *F* (*v, u*) stores the information of whether 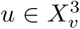. With 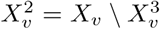, the updated workflow is illustrated in Figure 5.

**Figure 4:**
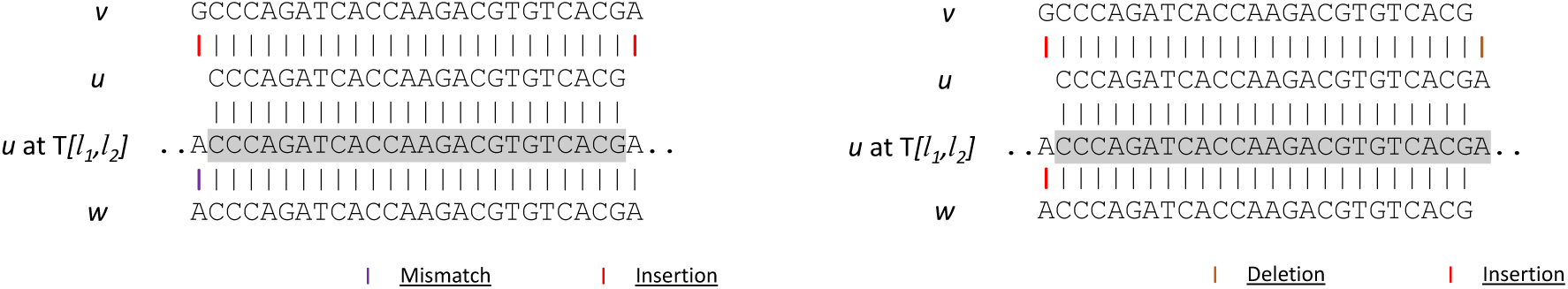
Illustration of Lemma 4. The figure to the left shows an example where *u* is a substring of *v*, while the figure to the right shows an example where *u* is an overlapping string of *v*. Notice that in both figures, *w* is always optimally aligned to *v*.

**Figure 5:**
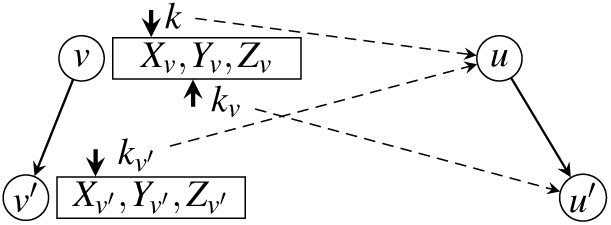
Illustration of adding *Z*_*v*_:= *{F* (*u, v*) *| u ∈ X*_*v*_*}* to each node.

Computation of *F* (*v, u*) can be piggybacked on top of computing *D*(*v, u*) in Algorithm 1. Given *u* and *v, F* (*v, u*) stores whether *v* and *u* possess any of the below *immediate conditions*:

(1) *v* is a prefix of *u*. (2) *v* is a suffix of *u*. (3) *u* is a prefix of *v*. (4) *u* is a suffix of *v*. (5) *v* is neither a prefix nor a suffix but a substring of *u*. (6) *u* is neither a prefix nor a suffix but a substring of *v*. (7) A prefix of *v* is a suffix of *u*. (8) A suffix of *v* is a prefix of *u*.

From above immediate conditions, we deduce the immediate relationship between *v* and *u*. With conditions 1–6, we can infer the superstring-substring relationship. With Condition 7–8, we can infer the overlapping relationship. If *v* and *u* qualifies none of the above immediate conditions, then they must be non-immediate neighbors.

For simplicity, we initialize each node as satisfying immediate conditions 1, 2, 3 and 4 to itself. We initialize the root node *r* as a prefix to any of its neighbors; and any neighbors of *r* as a suffix to *r*. Finally, *r* is not an overlapping string or a substring of any neighbor.

*F* (*u, v*) can be computed in constant time if *F* (*p*(*v*), *p*(*u*)), *F* (*p*(*v*), *u*) and *F* (*v, p*(*u*)) are known. For example, in Figure 5, *F* (*v′, u′*) satisfies condition 1, only if (a) *v′* = *u′* or (b) *F* (*v′, u*) satisfies condition 1. *F* (*v′, u′*) satisfies condition 2, only if (a) *u′* = *v′* or (b) *F* (*v, u*) satisfies condition 2 and *σ*(*u, u′*) = *σ*(*v, v′*). Conditions 3 and 4 are mirror cases of conditions 1 and 2, respectively with *v* and *u, v′* and *u′* trading places. *F* (*v′, u′*) satisfies condition 5, only if (a) *F* (*v′, u*) satisfies condition 5 or (b) *F* (*v′, u*) satisfies condition 2, while *v′* ≠ *u′* and *v* is not root. Condition 6 is a mirror case of condition 5. *F* (*v′, u′*) satisfies condition 7 only if (a) *F* (*v, u′*) satisfies condition 7 or (b) *F* (*v, u′*) satisfies condition 2, while *v* ≠ *u′* and *v* is not root. Condition 8 is a mirror case of condition 7.

The computation of *F* (·, ·) is piggybacked on top of the computation of *D*(·, ·), as both methods use dynamic programming. Both methods require piror knowledge between the child-parent and parent-parent node pairs; and from prior results, both methods compute the new result of the child-child node pair in constant time. As a result, piggybacking the computation of immediateness does not increase the complexity of Algorithm 1.

Finally, the confidence radius of node *v* equals min *D*(*v, u*) where 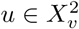, where *F* (*v, u*) does not satisfy any of the immediate conditions. The confidence radius of a node can be found by simply scanning its neighbor array, which finishes in linear time. The overall complexity of constructing the confidence radius database is still *O*(*|*Σ*|*^2^ · *M)*.

The confidence radius database is stored in a *| T |*-by-*P* table, where *P* is user-provided. The [*x, y*] entry of the table stores the *c*_*s*_ of seed *T* [*x, x* + *y*]. In practice, | *T*| *≫ P* and when needed, we can condense the confidence radius database into bit-vectors to reduce the table size. If necessary, when *| T |* is large, we can sub-sample seeds only at fixed-length intervals to further reduce the storage footprint.

## 4 A Seeding Scheme with Context-Aware Seeds

While the major goal of this paper is to establish the theoretical framework of CAS, to test the effectiveness of CAS, we propose a greedy seed selection method, referred to as greedy CAS seeding. Greedy CAS seeding selects consecutive Maximum Exact Matching substrings (MEMs, which are seeds that cannot be further extended without bumping into errors) from a read as seeds. At the end of each MEM, greedy CAS seeding heuristically skips the next two base pairs, in an effort to skip potential errors. Greedy CAS seeding sorts seeds by their frequency from low to high, into *S*_*raw*_. Then selects the minimum number of seeds *S* from *S*_*raw*_ in sequential order such that Σ_*s∈S*_ *c*_*s*_ *≥ t.*. In the rare cases where there is insufficient number of CAS seeds such that ∄*S* with Σ_*s∈S*_ *c*_*s*_ *≥ t,* greedy CAS seeding reverts back to using the pigeonhole principle, by dividing the read into *t* non-overlapping seeds.

Figure 6 compares the seed extraction results of greedy CAS seeding against the state-of-the-art, pigeonhole-principle-based seeding method, the Optimal Seed Solver (OSS). OSS has been previously shown that it generates the least frequent seeds, when compared to other pigeonhole-principle-based seeding methods, such as flexible-placement k-mers or spaced seeds. Figure 6 demonstrates both seeding methods in action under *t* = 4. Greedy CAS seeding is shown in the upper half while OSS is shown in the lower half. Compared to OSS, which uses a total of *t* = 4 seeds, greedy CAS seeding uses only two seeds. As a result, greedy CAS seeding can afford longer and less frequent seeds.

**Figure 6:**
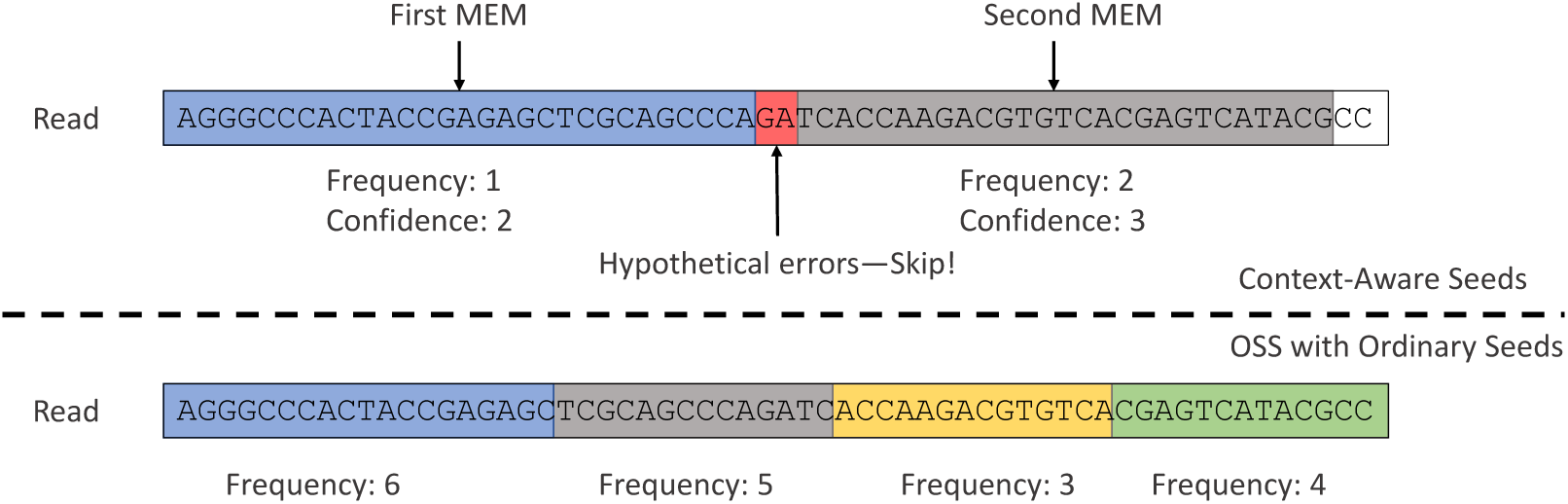
An example of drawing context-aware seeds from a read.

Greedy CAS seeding has a maximum complexity of *O*(*|R|* + *|S|* log(*|S|*)) (*|R|* denotes the length of *R* while *|S|* denotes the cardinality of set *S*). We use Burrows-Wheelers Transformation (BWT) array to index seeds. With BWT array, it takes *O*(*|s|*) operations to access the seed database for seed *s* and locate all seed locations of *s*. Given that Σ _*s∈S*_*|s| ≤ |R|*, and *|S| ≤ t ≪ R*, we conclude that the maximum complexity of greedy CAS seeding is *O*(*R* + *t* log(*t*)).

## 5 Experiments

We benchmark greedy CAS seeding against OSS on the *E. coli* genome. We benchmark both seeding schemes on a 22-million, 100-bp *E. coli* read set from EMBL-EBI, ERX008638-1. We build a confidence radius database for *E. coli* genome with a maximum edit distance threshold *t* = 5. We measure the effectiveness of both approaches by comparing the average total frequency of selected seeds under different edit distance thresholds *t* ={1, 2, 3, 4, 5}.

Figure 7 shows the average total seed frequency comparison between the two approaches. OSS has slightly smaller total seed frequency (averaged over all reads) under *t* = 1, but it quickly increases, exceeding CAS at *t >* 1. OSS out performs CAS under *t* = 1 because greedy CAS seeding extracts seeds sequentially; while OSS scans through all possible MEM placements in a read and picks the least frequent placement. When *t* gets larger, OSS is pressured to use more seeds, which leads to using shorter and more frequent seeds. To the contrary, greedy CAS seeding often uses fewer than *t* seeds, as shown in Figure 8, which let it use longer and less frequent seeds. At *t* = 4, greedy CAS seeding out performs OSS by 25.4%.

**Figure 7:**
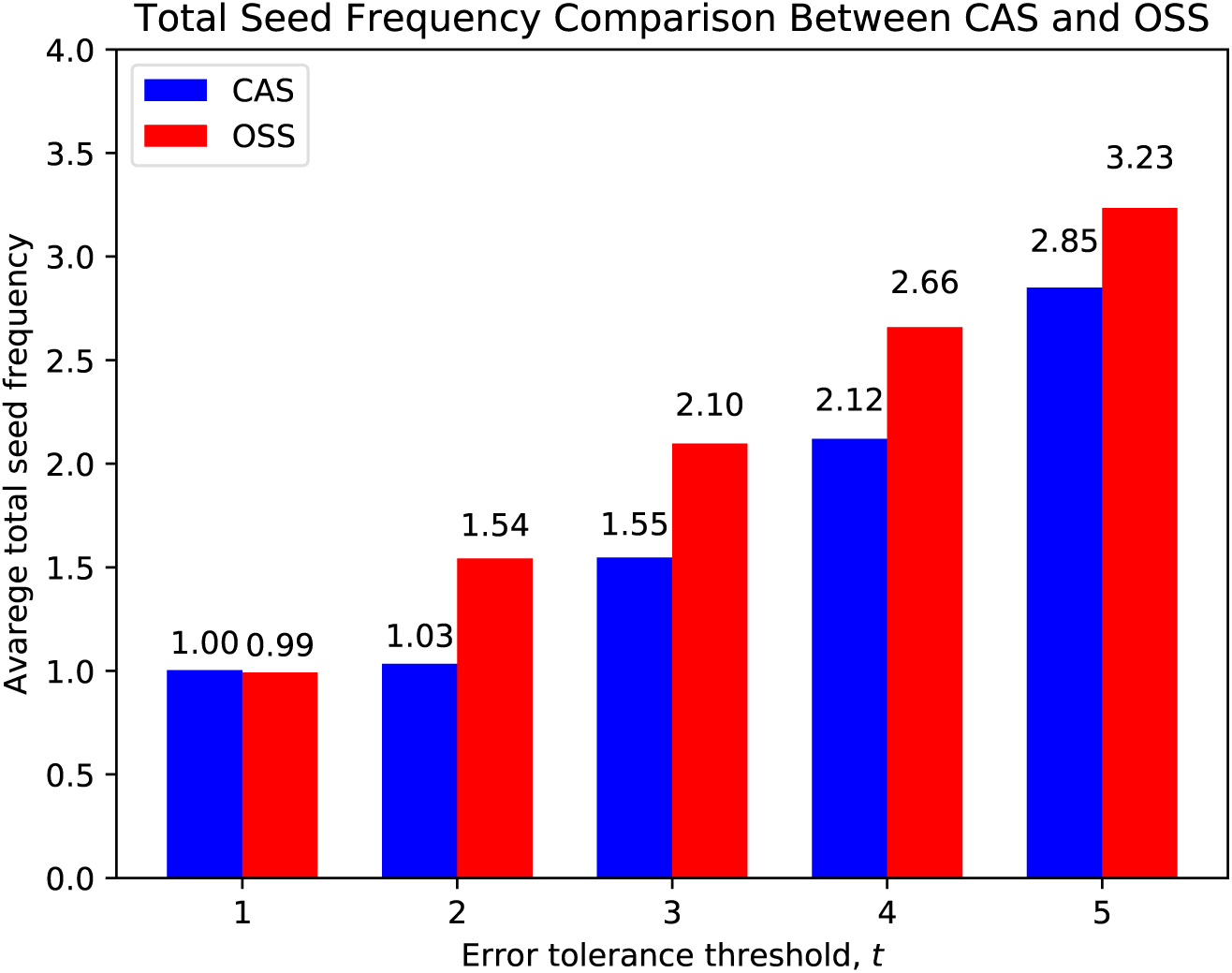
Comparison between CAS and OSS in terms of total seed frequency, with variate edit distance thresholds *t*.

**Figure 8:**
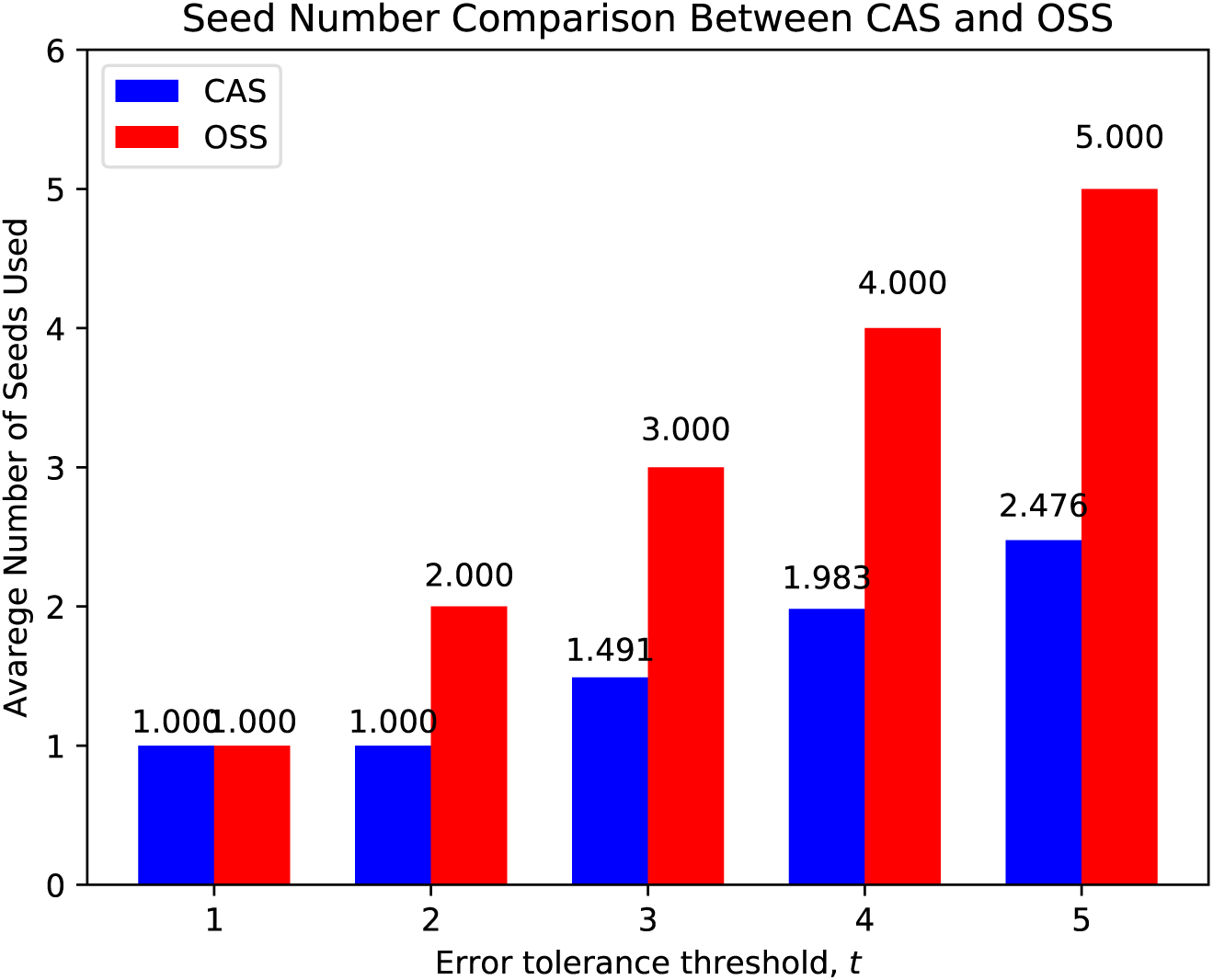
Comparison between CAS and OSS in terms of average number of seeds used, with variate edit distance thresholds *t*.

CAS is expected to perform better on larger genomes. The *E. coli* genome is a small genome, which has only around 4.6 million base pairs. In comparison, the human genome has more than 3 billion base pairs. For small genomes, seeds becomes less frequent by nature. Therefore short seeds becomes acceptable as they are not as frequent as they are in larger genomes. We expect CAS to perform better in larger genomes. However, due to practical (not theoretical) limitations in scaling up the construction of the confidence radius database on larger genomes (further elaborated in the Discussion section), we only demonstrate CAS on the *E. coli* genome.

While the focus of this paper is to establish the theoretical foundation of CAS, instead of providing a complete read mapping solution, it is worth mentioning that greedy CAS seeding (only the seeding mechanism) is more practical than OSS. OSS requires scanning through all substrings, denoted as set *S*_*OSS*_, of *R*, which has a total size of *O*(*|R|*^2^), for seed frequencies. Combined with BWT, it takes at least *O*(*|R|*^2^) operations to collect all seed frequencies with OSS. Greedy CAS seeding, to the contrary, finishes in *O*(*|R|* + *tlog*(*t*)) time with *t ≫ |R|*.

## 6 Discussion

Although Algorithm 1 finishes in *O*(*|*Σ*|*^2^ · *M)* time, in practice, *M* could be on the scale of trillions or more, for large and complex reference genomes. This is because for large genomes, the suffix trie is close to full in the first ten to twenty levels, where almost every permutation of letters exists. Nodes in these levels have large numbers of neighbors: the number of neighbors of a node *v*, equals to the number of unique strings formed by editing the string of *v* with up to *t* edits. After each edit, the resulting string is guaranteed to appear in *Trie*. This is further amplified by the exponentially-growing number of nodes in each level. In human genomes, there are more than one billion unique 15-base-pair suffixes. This means that for human genomes, under *t* = 4, there could be more than 1 trillion total neighbors just for 15-base-pair suffixes. Maintaining metadata at such scale vastly exceeds the capacity of our currently available computational power.

While there are many nodes (long suffixes) with fewer neighbors, given that Algorithm 1 traverses *Trie* in a top-down manner, it is unavoidable to track the massive number of neighbors for short suffixes. This is an interesting algorithmic problem for future work.

CAS may be applied to situations other than NGS read mapping. For example, the idea of context-aware seeds may improve long-read mapping. Long reads suffer from high error rates [3, 4, 13]. Finding error-free seeds for long reads is very challenging [5]. CAS can serve as a metric measuring the likelihood of seeds having errors: if there exists a seed, *s*, with high confidence radius, it is highly likely that *s* is free of errors. The likelihood of obtaining a reference-matching seed through many accidental errors is small.

Finally, CAS can be applied to develop probes for DNA and RNA identification. When designing probe sequences, it is important to make certain that the target sequence is unique in the genome [2, 10, 12]. It prevents probes from accidentally annealing to a similar sequences. CAS checks the existence of similar sequences by consulting the confidence radius database.

## 7 Conclusion

In this work, we proposed a new seeding framework, context-aware seeds (CAS). CAS extends the pigeonhole principle and guarantees finding all valid mappings with fewer seeds. CAS associates each seed *s* with a confidence radius *c*_*s*_, defined as a lower bound of edit distances towards nontrivial neighbors of *s*. We proved that the CAS can find all valid mappings of any read *R*, as long as its seeds *s* satisfy Σ*c*_*s*_ *≥ t*.

We proposed a linear-time algorithm for constructing the confidence radius database. It computes the confidence radii of seeds by traversing the suffix trie of a reference. We experimented CAS on *E. coli* genome and compared it against the state-of-the-art pigeonhole-principle-based seeding scheme, OSS, and showed that CAS outperforms OSS by reducing the sum of seed frequencies by up to 25.4%.

This paper focuses on the theoretical aspects of CAS, especially how it extends the pigeonhole principle into using fewer seeds. Composing a practical solution of Algorithm 1 on larger genomes is an interesting-yet-separate problem for future work.

## Acknowledgements

This research is funded in part by the Gordon and Betty Moore Foundations Data-Driven Discovery Initiative through grant GBMF4554 to CK, by the U.S. National Science Foundation (CCF-1319998) and by the U.S. National Institutes of Health (R01GM122935). This work was partially funded by the Shurl and Kay Curci Foundation. This project is funded, in part, by a grant (4100070287) from the Pennsylvania Department of Health. The department specifically disclaims responsibility for any analyses, interpretations, or conclusions.

